# The role of claw colouration in fiddler crabs’ mate choice: Private communication channel or handicap?

**DOI:** 10.1101/2021.02.06.430085

**Authors:** Diogo Jackson Aquino Silva, Marilia Fernandes Erickson, Raiane dos Santos Guidi, Daniel Marques Almeida Pessoa

## Abstract

Colour cues play an important role in sexual selection and conspecific recognition. Literature shows that conspecifics might enjoy their everyday chat, without ever worrying about occasional eavesdroppers (e.g., predators), when information interchange evolves into a private communication channel. Yet, when signalling is converted into foraging cues by predators, their prey must pay the due cost for sustaining conversation. For that matter, fiddler crabs draw attention for having flashy enlarged claws that could potentially attract the attention of many predators. Surprisingly, the costs associated with claw colouration in fiddler crabs are still poorly understood and have never been studied in American species. Here, we initially examine whether hypertrophied claws of American thin-fingered fiddler crabs (*Leptuca leptodactyla*) reflect UV-light and how conspecific females react to these cues. Then we test two alternative hypotheses concerning the role of claw colouration in fiddler crabs’ mate choice: a) that claw colouration evolved into a private communication channel, which could have significantly lowered signalling costs for males; b) that claw colouration is conspicuous to potential reproductive partners, as well as to predators, making colour signalling by males very costly (i.e., a handicap). Thereafter, we measured the reflectance spectra from several enlarged claws and modelled their chromatic contrast against the background spectrum, considering the visual systems of conspecific fiddler crabs and two kinds of predators (terns and plovers). We also tested female conspecifics’ preference towards enlarged claws that reflected UV-light or other colour cues, by artificially altering claw colouration. Our results show a clear female preference for UV reflecting males. We also found that natural enlarged claws should be highly detectable by avian predators, refuting the private communication channel hypothesis. Furthermore, since female fiddler crabs select the most flamboyant claws from the sandy background, claw colouration in fiddler crabs can be understood as an honest signal.

**HIGHLIGHTS:**

1. The hypertrophied claws of male *Leptuca leptodactyla* reflect UV light.
2. Female fiddler crabs display a natural preference for UV light cues.
3. Conspicuous claws function as handicaps and may honestly signal individual quality.
4. Male enlarged claws are more conspicuous to birds than to crabs.
5. Our data refute the presence of a private communication channel in *L. leptodactyla*.

## INTRODUCTION

The evolution of animal colouration has been associated with numerous functional drivers, such as social signalling, antipredator defence, parasitic exploitation, thermoregulation, and protective colouration (Cuthill et al. 2017). Among these drivers, the use of colour signals for intra and interspecific recognition, intrasexual competition, and mate choice have received special attention from researchers that study different animal taxa (Higham and Winter 2015). Yet, when we consider the study of the functional significance of colouration, an especially colourful group that has received far less attention, at least when compared to other arthropods (e.g., insects and spiders), is Brachyura (crabs) (Caro 2018), a promising taxon on which to test some key evolutionary drivers of external appearance (Zeil and Hemmi 2006; Caro 2018).

Among Brachyura, fiddler crabs have attracted the scientific interest for some time, and, for that matter, it is a group that already offers a substantial theoretical framework on which working hypotheses may be formulated. To wit, literature has established these animals present an array of agonistic and sexual visual displays based on size (Oliveira and Custodio 1998), posture (Schöne 1968; Murai and Backwell 2006), movement (Byers et al. 2010) and colour (Detto 2007) cues. Such visual cues are thought to relay information concerning individual reproductive status (Crane 1975) and quality (Latruffe et al. 1999), and to allow intra and interspecific recognition (Dyson et al. 2020).

Typically, male fiddler crabs have a brightly coloured (Dyson and Backwell 2016) and hypertrophied claw that can be used either as a weapon, in male-male disputes, or as an ornament, for female attraction (Crane 1975). Since fiddler crabs express two different kinds of visual pigments, which are responsible for absorbing light in the UV (300-400 nm) and visible (400-700 nm) light range (Horch et al. 2002; Jordão et al. 2007), they are expected to see UV cues and to show a dichromatic colour discrimination (Jacobs 2018). Behavioural experiments, conducted on banana fiddler crabs (*Austruca mjoebergi*), suggest that colour (UV light included), but not brightness, is important for conspecific reckoning and mate choice (Detto et al. 2006; Detto 2007; Detto and Backwell 2009; Dyson et al. 2020).

The adaptive value of UV light reflection in fiddler crabs, however, is still poorly understood and might be attributed to different ecological interactions. For instance, Detto and Backwell (2009) hypothesize that, if UV reflection enhances male conspicuity, increasing their detectability by female fiddler crabs, the trait should have been fixed in the population. On one hand, the argument of Detto and Backwell (2009) seems to follow Fisher’s runaway selection hypothesis (see Henshaw and Jones 2020), according to which a secondary sexual trait expressed in one sex should become correlated with a preference for the trait in the other sex. On the other hand, UV reflection could also be understood as a handicap that attracts the attention of predators and warrants signal honesty about individual quality to reproductive partners (Zahavi 1975). Bright colourations (which also include UV reflection) has been pointed out as honest indicators of low parasite loads (Hamilton and Zuk 1982) or, in a more general sense, good genes (Andersson et al. 1998). Yet, these possibilities remain to be tested in fiddler crabs.

Another explanation is that ultraviolet reflection could also result from a trade-off between the advantages of intraspecific conspicuous signalling and the disadvantages of predator attraction (Hemmi et al. 2006; Cummings et al. 2008). Fish subjected to these conditions have been reported to develop UV light reflection, which could yield a private communication channel (i.e., wherein a species produces a signal that is detected by conspecifics but not predators; see Cummings et al. 2003). In fiddler crabs’ list of predators, we usually find many bird species, which detect UV light very well, such as plovers (Ribeiro et al. 2003) and terns (Land 1999).

Recently, a taxonomic review of family Ocypodidae Rafinesque, 1815 (Crustacea: Brachyura) divided fiddler crabs’ former taxon (i.e., *Uca*) in 11 different genera (Shih et al. 2016). The endemic genera from the Americas (*Leptuca*, *Minuca*, *Petruca* and *Uca*) now encompass 57 species, approximately 55% of fiddler crabs’ current species. Among all 104 fiddler crab species, only four (*Afruca tangeri*, *Austruca mjoebergi*, *Tubuca signata* and *Tubuca capricornis*), from Europe/Africa and Australia, already had the utility of their claw colouration examined (Cummings et al. 2008; Detto et al. 2006; Detto 2007; Detto and Backwell 2009; Dyson et al. 2020). More surprisingly, *Leptuca*, fiddler crabs’ most heavily studied taxon (Nabout et al. 2010), and the richest genus in Ocypodidae family, enclosing almost one third of all fiddler crab species (i.e., 30 species according to Shih et al. 2016), has not yet been explored with respect to the ecological pressures shaping its claw colouration.

Hence, at first, we test two competing hypotheses regarding the functional significance of fiddler crabs’ enlarged claw colour, using *Leptuca leptodactyla* as an experimental model. (I) We hypothesize that hypertrophied claw colouration plays a part in a private communication channel, through which fiddler crabs exchange social/reproductive signals that are not detectable by their predators. We predict our results will show that colour contrast between enlarged claws and the sandy background is significantly perceptible to fiddler crabs’ visual system, while imperceptible to the visual systems of their potential predators. (II) As an alternative hypothesis, we propose that the conspicuity of enlarged claws works as a handicap (see Zahavi 1975), imposing to male fiddler crabs the heavy cost of enhancing predator attraction, while also honestly advertising their presence and quality to potential female mates. In this case, we predict our results will show the more noticeable an enlarged claw is for a reproductive partner, the more conspicuous it should also be for predators.

In addition, we also test a third hypothesis regarding the role that UV light reflectance exerts on female mate choice. (III) We hypothesize that UV reflection from enlarged claws will be the major colour signal considered by females in their decision making. In case our third hypothesis is correct, we expect our behavioural results to demonstrate females prefer males that reflect UV light, as already shown for Australian species (Detto and Backwell 2009).

## METHODS

### Choice of Study Species and Study Site

Just a few reports have examined the biology of the thin-fingered fiddler crabs (*Leptuca leptodactyla*; Figure 1) (Nabout et al. 2010), which populations spread throughout the Western Atlantic: Caribbean, Venezuela to Brazil, eastern Yucatan Peninsula (Mexico), south-eastern Florida (USA). The present study took place at a mangrove area, in which a population of thin-fingered fiddler crabs (*L. leptodactyla*) naturally occurs, showing a rainy tropical climate, with rains extending from February to September, and temperatures ranging from 21°C (min.) to 30°C (max.), with a mean temperature of 26°C, and predominant vegetation comprised of sandy coastal plains and mangroves. The area is located within a hydrographic basin, in which unconsolidated sand and gravel composes the fluvial flatland, that is subject to periodical flooding. At the study site, substrate becomes exposed during the low tides, revealing hundreds of burrows, in which *L. leptodactyla* and other crab species live.

**Figure 1.**
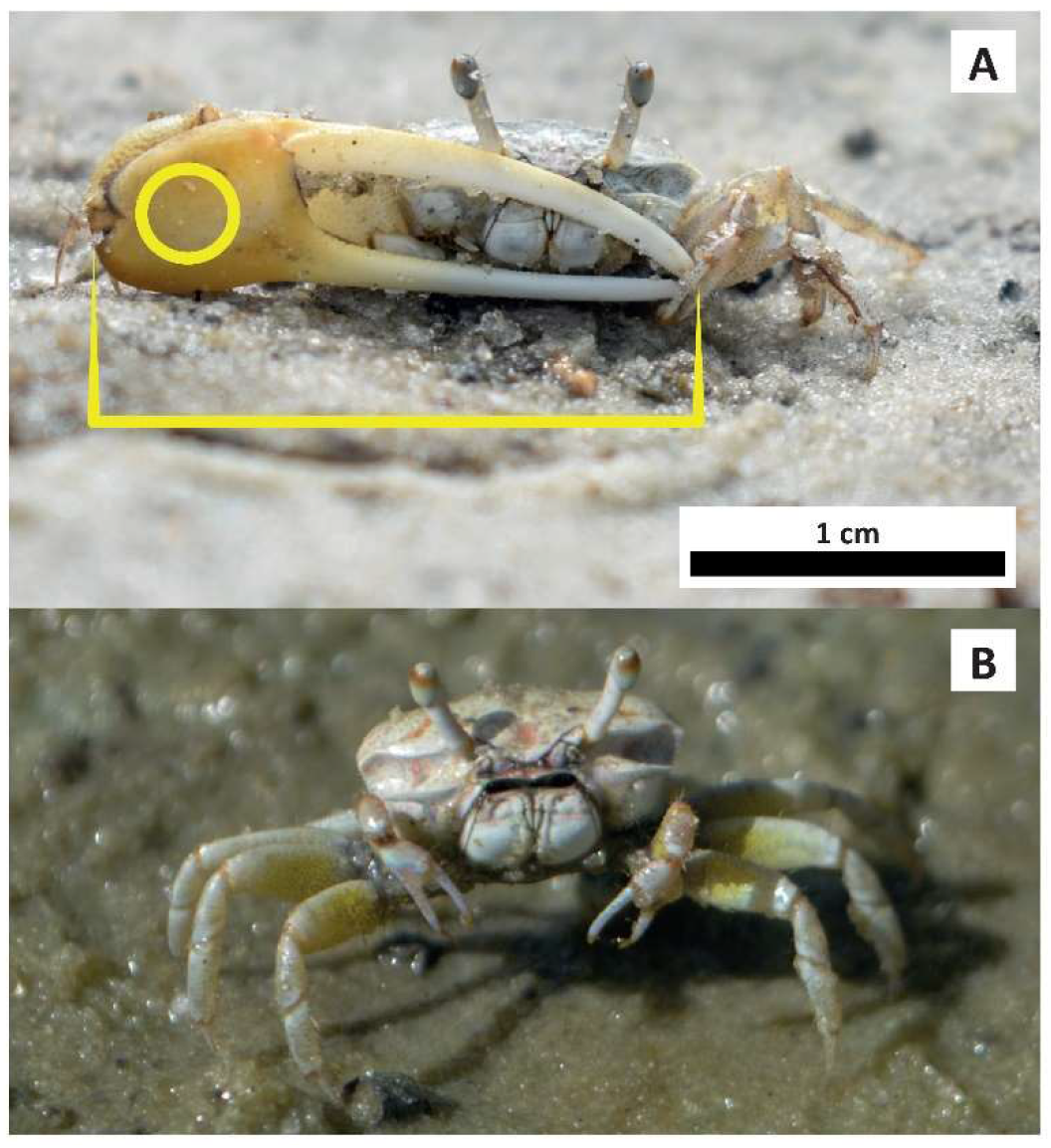
Male (A) and female (B) specimens of thin-fingered fiddler crabs (*Leptuca leptodactyla*). Yellow callipers indicate how male hypertrophied claws were sized, while the yellow circle indicates where the reflectance spectra (colour) of male hypertrophied claws were measured.

### Mate Choice Experiments

We conducted experiments throughout the low tides, and our experimental procedures were adapted from Detto et al. (2006), Detto (2007) and Detto and Backwell (2009). Before the beginning of each experimental session, we delimited a squared shaped arena (30 cm²) on the same sandy substrate in which the animals built their burrows, foraged and mate. For that, we displaced all the animals, obstructed their burrows, and drew a square in the sand.

Then, we captured a few male fiddler crabs (*L. leptodactyla*) and measured the length (Figure 1) of their hypertrophied claws (chelae) with a calliper, from the base of their palm (propodus) to their fingertip (dactyl), assigning the crabs to different groups according to their claw’s size. The maximal acceptable intragroup difference was established at 0.1 cm. From a same group, we randomly chose a set of four size-matched male crabs and subjected each animal to one of the experimental treatments described in Table 1. We decided to use white, yellow, and orange paints because white is *L. leptodactyla*’s carapace colour, while different shades of yellow and orange are the most frequent colouration found on their hypertrophied claws. We also decided to use blue paint to represent a kind of unfamiliar colour that is not expressed by *L. leptodactyla*, or any other species found in the same crab community, but that can be found on species of fiddler crabs of other geographical regions. In experiment 1, irrespective of their experimental treatment, males’ posterior carapaces were also treated with white paint and/or sunblock (Table 1), to control for any chemical cues that could be transmitted to the females. Males’ posterior carapaces were not viewable from females’ location.

**Table 1.**
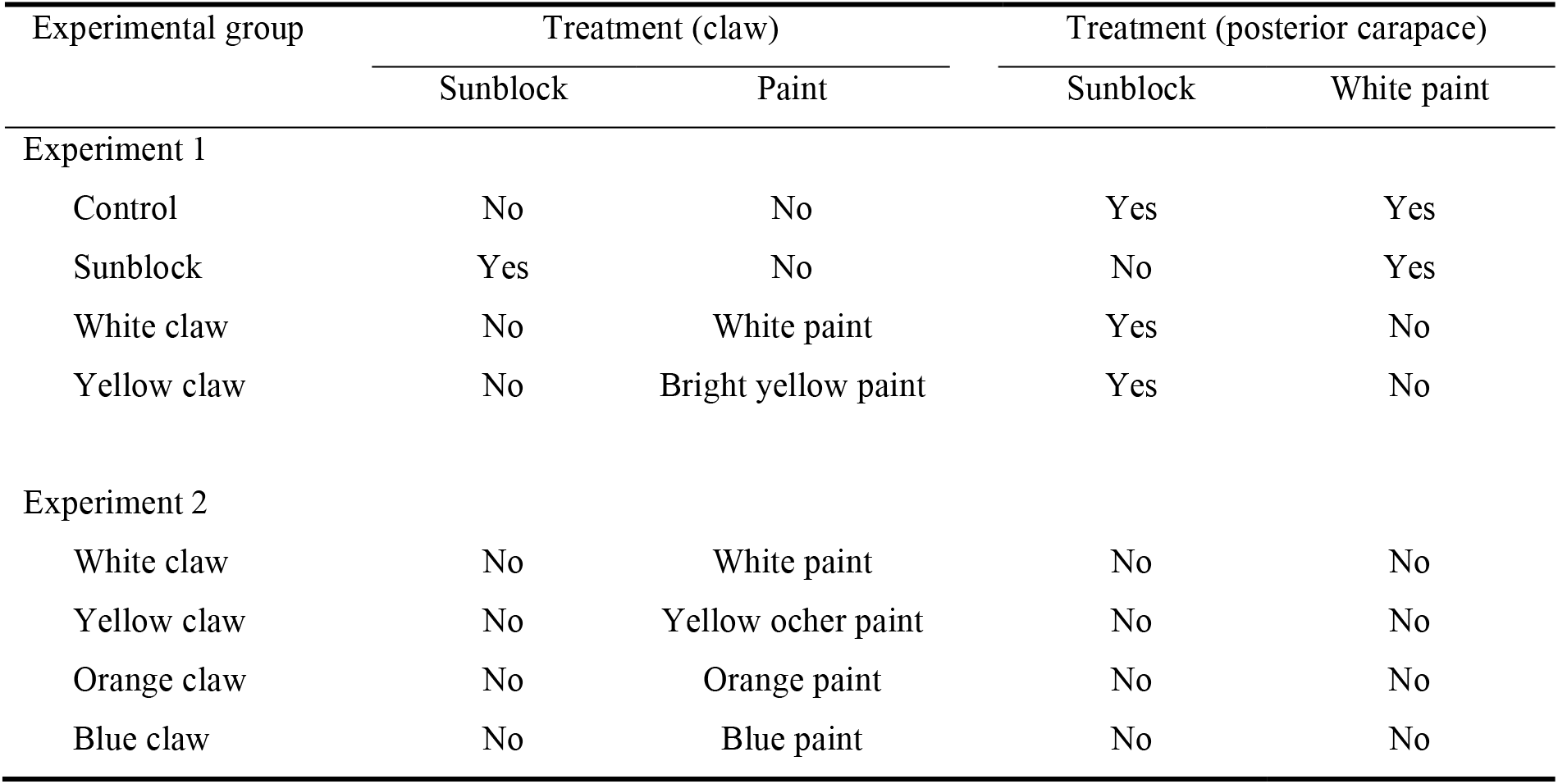
Treatments to which male thin-fingered fiddler crabs (*Leptuca leptodactyla*) were assigned in experiments 1 and 2. All paints were acquired from Acrilex (Matte Ink for Crafts) and sunblock was acquired from Natura (Fotoequilibrio SPF 60).

| Experimental group | Treatment (claw) |  | Treatment (posterior carapace) |  |
| --- | --- | --- | --- | --- |
|  | Sunblock | Paint | Sunblock | White paint |
| Experiment 1 |  |  |  |  |
| Control | No | No | Yes | Yes |
| Sunblock | Yes | No | No | Yes |
| White claw | No | White paint | Yes | No |
| Yellow claw | No | Bright yellow paint | Yes | No |
| Experiment 2 |  |  |  |  |
| White claw | No | White paint | No | No |
| Yellow claw | No | Yellow ocher paint | No | No |
| Orange claw | No | Orange paint | No | No |
| Blue claw | No | Blue paint | No | No |

Once painted by the experimenters, males were attached to nylon threads through their posterior carapaces, with superglue, while the other end of the threads were tied to nails, anchoring the crabs to the arena’s substrate, one male crab at each corner, facing each other, at 12 cm from the centre. Anchoring prevented males from moving, but still allowed them to move their locomotory appendages and to wave their claws, although waving displays following capture were not observed. We always randomized treatment positioning between different sets of size-matched males.

Consecutively, we captured a female crab (*L. leptodactyla*), placed it at the centre of the arena, under a transparent cup, and kept it there, habituating, for one minute. After habituation, we lifted the cup and observed the female’s behaviour for three minutes, unless it spent less time making a valid choice (i.e., slowly approaching a male by less than two centimetres) or evading the arena. We considered females to have evaded the arena when, as soon as the cup was lifted, they quickly ran towards one of the males and left the arena, or they left the arena without approaching any male. Females that spent three minutes without making a valid choice or evading the arena were replaced by another one. Females were tested only once, being freed right away, however, each set of four size-matched males was presented to several females, until we recorded three female valid choices. After that, the set of males was released and replaced by a new group of four sized-matched males, and so on. Prior to their liberation, males were released from their nylon threads.

In experiment 1, we tested if females choose male conspecifics according to the presence of UV reflection. While in experiment 2, we tested if females showed any sensory bias towards any specific claw colouration (e.g., acquainted, and unacquainted colours), irrespective of UV cues. In experiment 1 we used 20 groups of males (totalling 80 males) and a total of 206 females (146 evasions and 60 female choices), while in experiment 2 we used 20 groups of males (80 males), having tested a total of 168 females (108 female evasions and 60 female choices). These sample sizes are in accordance with what has been stablished by previous published studies (Detto et al. 2006; Detto 2007; Detto and Backwell 2009).

Our research protocol adheres to the ASAB/ABS guidelines for the use of animals in research and is in accordance with institutional guidelines and local legislation.

### Spectral Measurements

We used a USB4000-UV-VIS (Ocean Optics, Inc.) spectrophotometer connected to a light source (DH2000-BAL; Ocean Optics, Inc.) through a bifurcated optic probe (QR450-7-XSR; Ocean Optics, Inc.). The tip of the optic probe was coupled to a custom-made probe holder, to reduce the sampling area (1 mm of diameter; Appendix 1). The system was calibrated using a WS-1-SL (Ocean Optics, Inc.), as the white standard, and by turning the light source off, as the black standard. All measurements were taken at a fixed angle (90º) and distance (5 mm) from stimuli, with help of the SpectraSuite software (Ocean Optics, Inc.), in which boxcar width and number of spectra averaged were set to 5 and 10, respectively.

Forty male fiddler crabs were, randomly, collected in the study area and brought to the laboratory, to have the natural colour of their hypertrophied claws registered (Figure 1). These data confirmed that the chosen population reflected UV light, and stablished that there was a natural variation among hypertrophied claw spectra. Using a shovel, we collected a sample of the study site’s sediment and carefully caried it to the laboratory to avoid disaggregation. Using the above-mentioned procedure, we also measured the reflectance spectrum of the outmost sediment layer (Figure 2).

**Figure 2.**
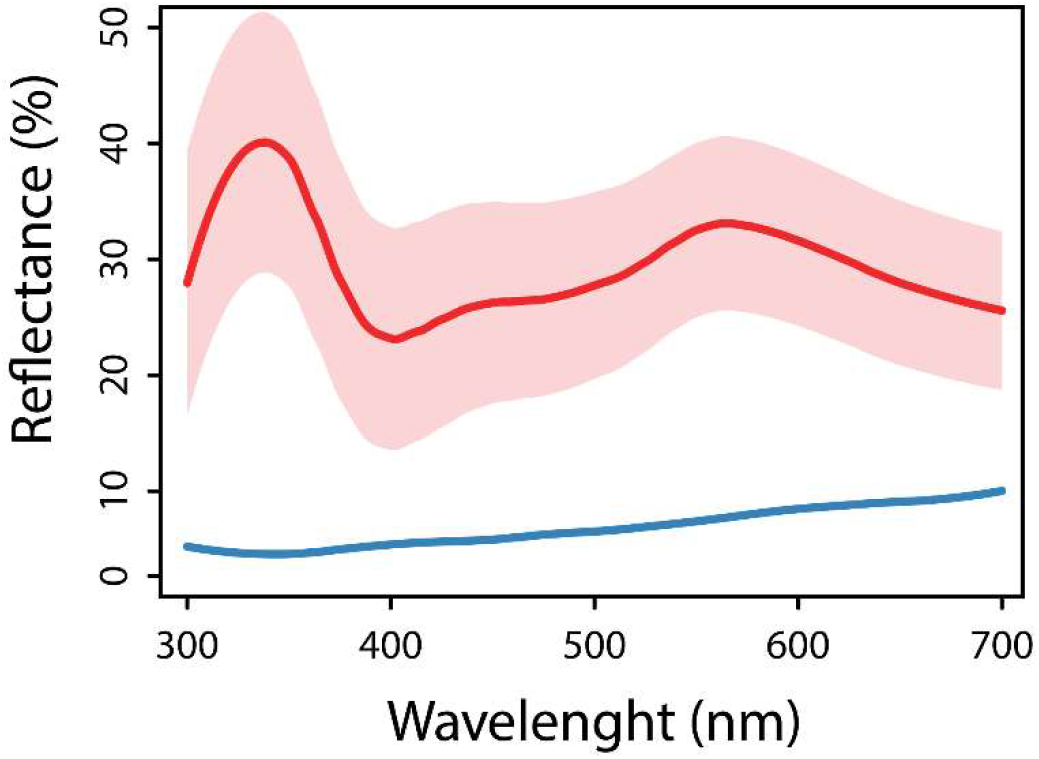
Natural reflectance spectra measured from a set of 40 male thin-fingered fiddler crabs’ (*Leptuca leptodactyla*) hypertrophied claws. Average spectrum and variation (maximal and minimum values) are indicated by the red line and its adjoining shaded pink area, respectively. The blue line represents the sandy background reflectance spectrum.

In a subsequent opportunity, in order to characterize the effect that each experimental treatment (Table 1) had on the reflectance spectra of male claws, we captured six additional crabs, measured their natural colouration, as described previously, and covered their hypertrophied claws with one of the following products: 1) sunblock (Natura Fotoequilibrio SPF 60); 2) white paint (Matte Ink for Crafts, Acrilex); 3) bright yellow paint (Matte Ink for Crafts, Acrilex); 4) yellow ochre paint (Matte Ink for Crafts, Acrilex); 5) orange paint (Matte Ink for Crafts, Acrilex); 6) blue paint (Matte Ink for Crafts, Acrilex). Males’ claw colourations were remeasured after treatment (Figure 3). All paints completely blocked UV light reflectance, in the same way sunblock did.

**Figure 3.**
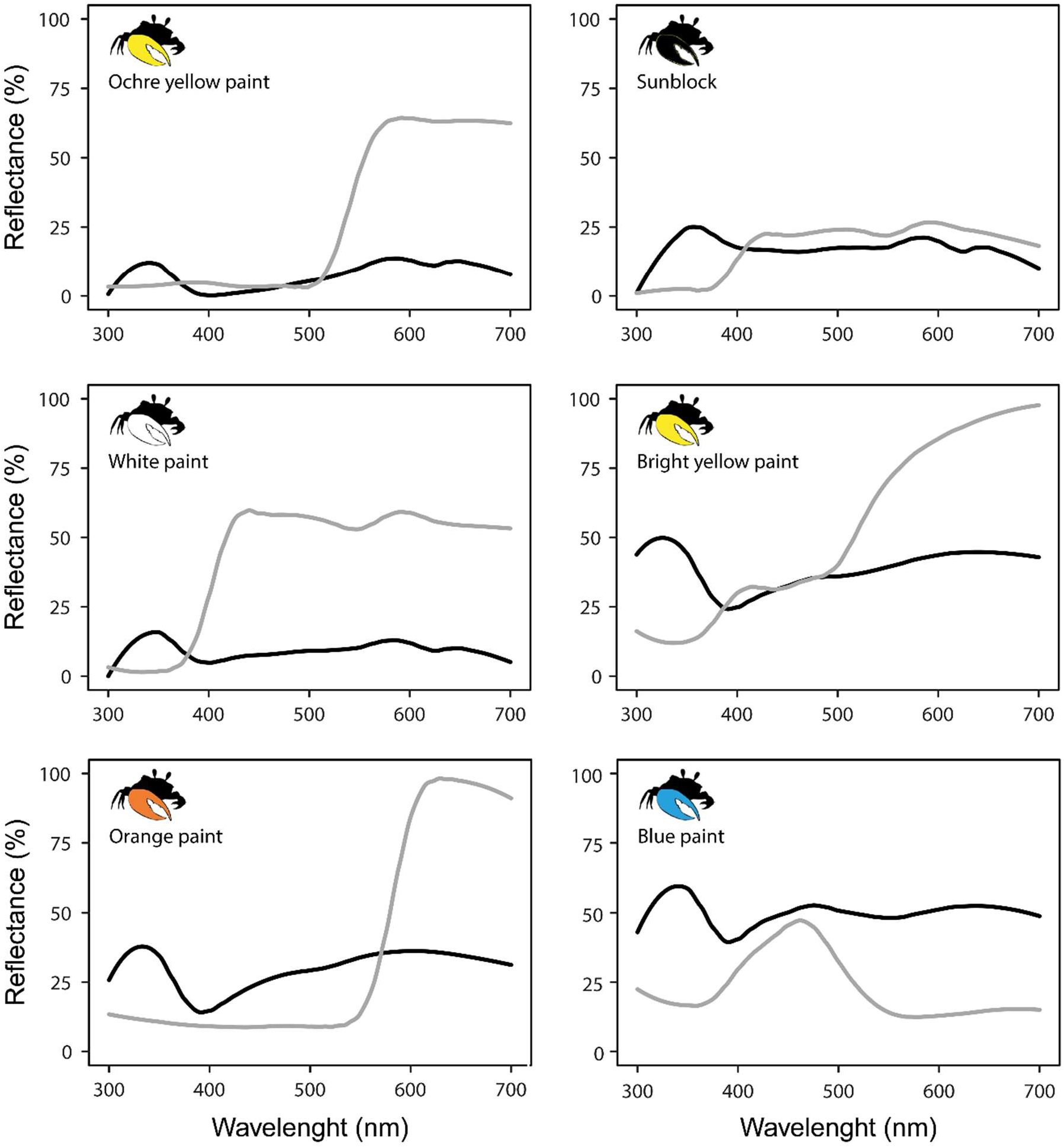
Reflectance spectra measured from the enlarged claws of six male thin-fingered fiddler crabs (*Leptuca leptodactyla*) subjected to different experimental treatments (detailed in Table 1). Natural spectra (before treatment) are represented by black lines, while grey lines indicate claws’ spectra after treatments.

### Visual Modelling

To determine how fiddler crabs and their predators (e.g., terns and plovers) visualize the colouration of male claws against the sandy substrate, we applied the receptor noise (RNL) model of colour discrimination (Vorobyev and Osorio 1998) to determine the chromatic contrasts (ΔS) between the reflectance spectrum of the sandy background and claws’ natural colourations, according to the visual system of fiddler crabs and birds. It is important to note that the RNL model only considers chromatic information, disregarding achromatic cues. Modelling was conducted in pavo 2.0 (Maia et al. 2019), a R package for spectral analysis of colour.

Initially, we estimated the absolute amount of light captured by the photoreceptors of each observer (‘Q_i_’ – quantum catch), considering three factors: 1) the reflectance spectrum of the object (i.e., male claw) or background (i.e., sandy substrate); 2) the illuminant spectrum of incident light (i.e., natural light shining on the study site); 3) the observer’s visual sensitivity curves. We compared the natural reflectance spectra collected from 40 animals, as described previously, with the reflectance spectrum collected from the sandy background (Figure 2). A standard daylight illuminant spectrum (i.e., illum = ‘D65’), provided by pavo 2.0’s library, was used. Since estimation of receptor sensitivities do not affect model results too seriously (Olsson et al. 2018), and because birds show two general colour vision phenotypes, those containing ultraviolet sensitive (UVS) cone photoreceptors and others containing violet sensitive (VS) cone photoreceptors, we used available spectral sensitivities from the Atlantic Mangrove Fiddler Crab (*Leptuca thayeri*: 430 nm, 520 nm; Horch et al. 2002), the Blue Tit (*Cyanistes caeruleus*: 372 nm, 453 nm, 539 nm, 607 nm; Hart and Hunt 2007) and the Peafowl (*Pavo cristatus*: 424 nm, 479 nm, 539 nm 607 nm; Hart and Hunt 2007), as proxies for fiddler crab’s, tern’s, and plover’s visual systems, respectively. For the visual system of crabs we set *trans* = ‘ideal’, while for terns we used *trans* = ‘bluetit’ (Hart et al. 2000). For plovers, we supplied *trans* as vector containing the ocular transmission spectra of *Pavo cristatus* (Hart 2002). Since we required absolute quantum catches, instead of relative quantum catches, we set *relative* = FALSE. We set the remaining parameters to default (i.e., vonkries = FALSE, scale = 1).

In addition, when calculating the chromatic contrasts (ΔS) for each kind of observer, we ran the RNL model and compared Q_i_ information of each individual crab with that of the sandy background, setting parameters to default (i.e., photoreceptor noise set as ‘neural’, weber.ref = ‘longest’) and including specific cone densities and photoreceptor noise values (i.e., weber fractions) for the visual systems of each modelled observer. For tern’s visual system we employed [n = c(1:1.9:2.7:2.7); based on *Cyanistes caeruleus* (Hart et al. 2000)], and for plover’s visual system we used [n = c(1:1.9:2.2:2.1); based on *Pavo cristatus* (Håstad et al. 2005)], applying pekin robin’s (*Leiothrix lutea*) weber = 0.1 (Maier and Bowmaker 1993) to both avian visual systems. Following previous publications (Hemmi et al. 2006; Cummings et al. 2008), for modelling fiddler crab’s visual system we applied honeybee’s (*Apis mellifera*) weber = 0.12 (Vorobyev et al. 2001) and used n = c(1:1), since no information about the proportion of photoreceptors is mentioned by literature. The ΔS output, between an object an its background, was given in units of just noticeable difference (JND). Following Siddiqi et al., (2004), we adopted three levels of detectability for the observers’ visual systems modelled in this study: cryptic (ΔS < 1 JND), detectable (1 JND ≤ ΔS ≤ 3 JND) and highly detectable (ΔS > 3 JND). The higher the chromatic contrast, the higher the colour difference between a male claw and the sandy background, favouring their detectability.

### Statistics

Owing to the non-parametric nature of our visual modelling results and behavioural data (Shapiro-Wilk, *P* < 0.05), chromatic contrast values generated by our visual model for different observers (i.e. crabs, terns and plovers) were compared through Kruskal-Wallis tests, with Dunn’s post hoc test and Bonferroni correction, while a Generalized linear model (Poisson model GZLM using a log link) was used to compare the different treatments of experiments 1 and 2. The Poisson model was described by ratio rate (RR) and confidence interval (CI). All analyses were run on R statistics (R Core Team, 2017), and assumed α = 0.05.

## RESULTS

### Visual Modelling

The output of our visual model (Figure 4), in which the natural colouration of 40 males was compared to the sandy substrate’s colour, shows that claws of *L. leptodactyla* produce colour signals that vary from cryptic (ΔS < 1) to highly detectable (ΔS > 3 JND), according to the eye of the beholder. When taking colour cues into consideration, most measured enlarged claws were expected to be conspicuous against the sand when seen by terns (i.e., 39 conspicuous claws out of 40) and by the plovers (i.e., 31 conspicuous claws out of 40), since most data points have fallen in the in the lighter grey area (i.e., detectable) or in the white area (i.e., highly detectable) of Figure 4.

**Figure 4.**
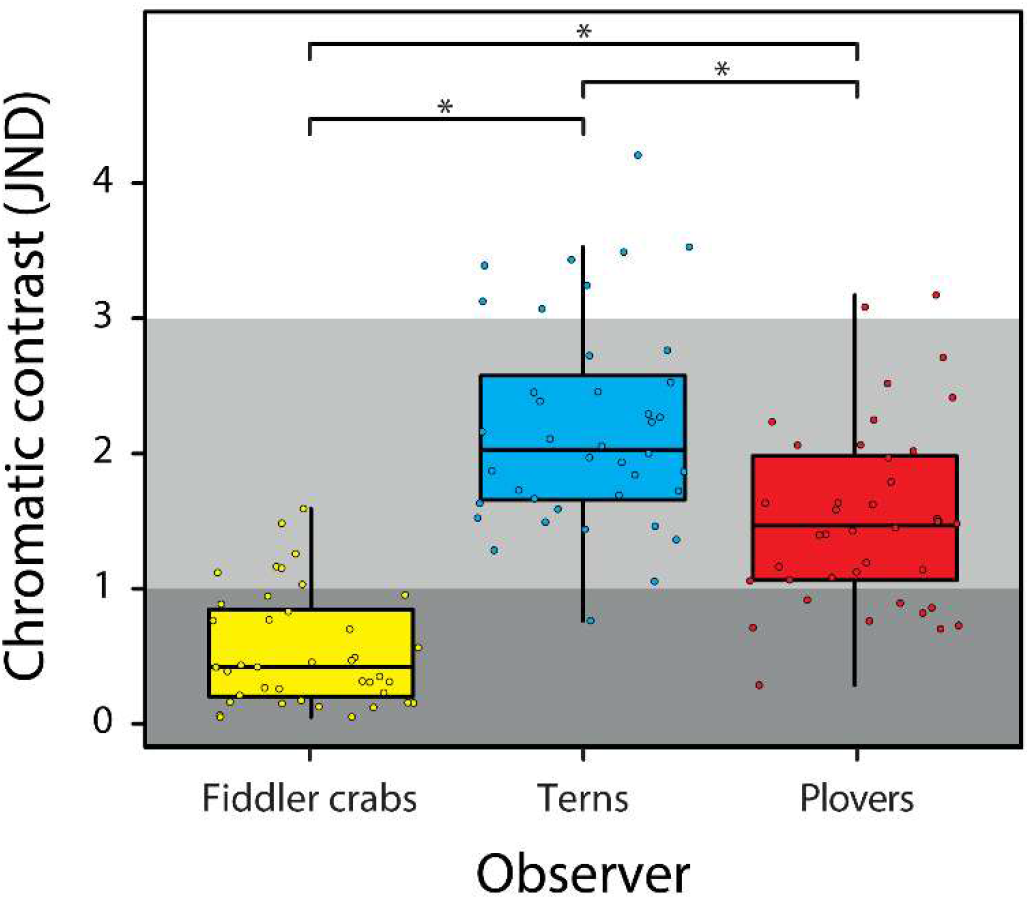
Chromatic contrast (ΔS) between the natural reflectance spectra measured from the hypertrophied claws of 40 male thin-fingered fiddler crabs (*Leptuca leptodactyla*) and the sandy substrate’s reflectance spectrum, modelled according to the visual systems of different observers. Dots represent individual values of chromatic contrast (in unit s of JND). Medians (black thick horizontal lines), interquartile ranges (boxes) and variability outside the upper and lower quartiles (whiskers) are also indicated. Thresholds of detectability are signalled by different grey areas. Dots that fall in the darker grey area are not detectable (ΔS < 1 JND), while dots that fall in the lighter grey area or white area are, respectively, detectable (1 JND ≤ ΔS ≤ 3 JND) and highly detectable (ΔS > 3 JND). Bars with asterisks indicate statistically significant difference between observers.

In contrast, for fiddler crabs’ conspecifics (e.g., females) only a minority (i.e., 7 out of 40) of enlarged claws have fallen above the darker grey area of Figure 4, which indicates that 33 males, out of 40, should not be able to attract the attention of females based on the colour of their claws alone. Seven out of 40 enlarged claws (17.5%), however, have fallen in the lighter grey area of detectability, while no claws were able to reach the white area of high detectability.

When we statistically compared chromatic contrast values generated by our visual model there was a significant difference in perceptual performance between different observers (Kruskal-Wallis: *χ^2^(2)* = 70.376; *P* < 0.0001; Figure 4), in which birds statistically outperform fiddler crabs (Dunn’s test: *P* < 0.0001), while terns outperform plovers (Dunn’s test: *P* < 0.001).

### Mate Choice Experiments

In Experiment 1, the Poisson model (GZLM) indicated that claw colouration influenced females preference (*χ^2^* = 12.928; *P* = 0.004; Figure 5a), favouring males showing UV light reflection (i.e. males with claws of natural colouration) when compared to males that had their UV light reflection depleted by sunblock (*RR* = 0.2962 [0.1 - 0.6]; *P* = 0.002), white paint (*RR* = 0.4074 [0.19 - 0.79]; *P* = 0.012), or yellow paint (RR = 0.5185 [0.26 - 0.97]; *P* = 0.046). While in Experiment 2, the Poisson model (GZLM) revealed no indication of a female sensory bias (*χ^2^* = 3.9329; *P* = 0.2688; Figure 5b) being directed towards any specific claw colouration, when UV light reflection was depleted in all available treatments.

**Figure 5.**
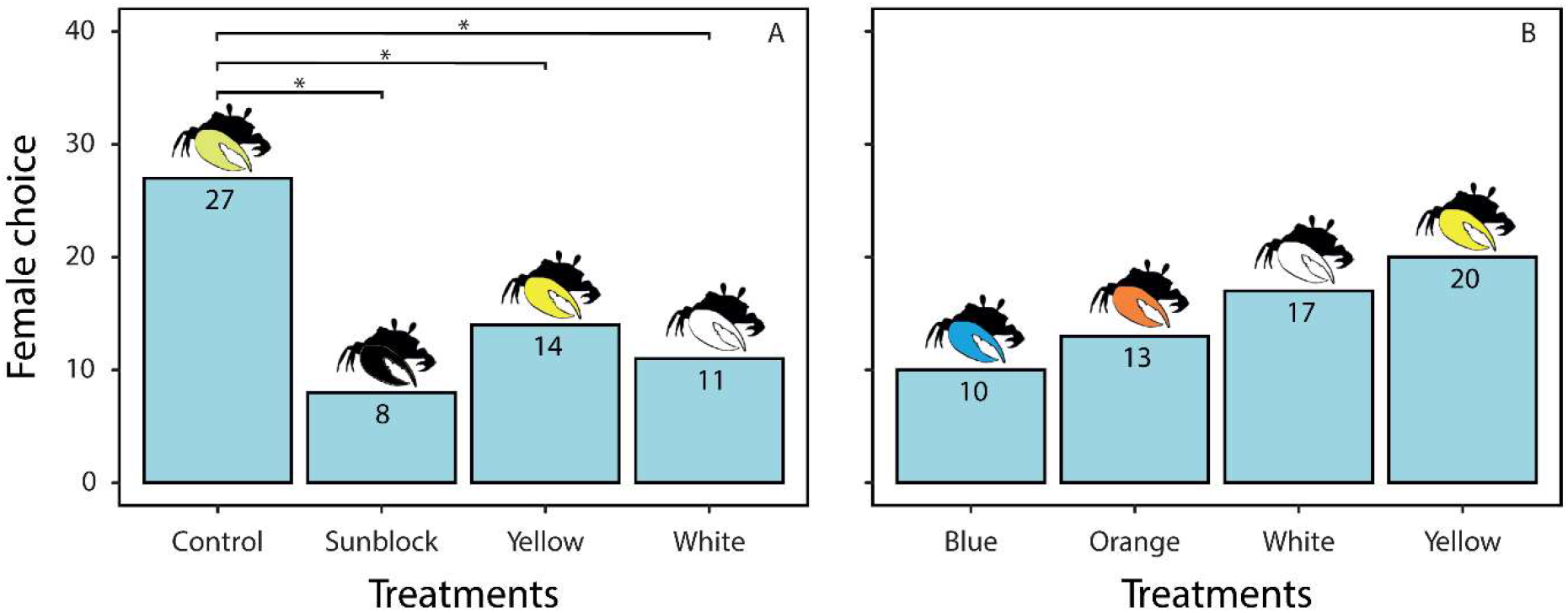
Mate choice by female thin-fingered fiddler crabs (*Leptuca leptodactyla*), when subjected to conditions of experiments 1 (A) and 2 (B). Treatments’ details are given in Table 1. Bars with asterisks indicate statistically significant difference between treatments.

## DISCUSSION

Our spectrophotometric data confirmed that the hypertrophied cheliped of thin-fingered fiddler crabs (*Leptuca leptodactyla*) significantly reflects UV light, while our behavioural results have proven that *L. leptodactyla* is also capable of discriminating the UV cues generated by conspecifics, which corroborates electrophysiological data gathered in a close species (*Leptuca thayeri* - Horch, 2002). Our findings parallel what has been found in an Australian species, the banana fiddler crab (*Austruca mjoebergi*), to which UV light reflection and preference has been associated with Fisherian explanations (Detto and Backwell 2009).

Yet, concerning non-UV cues, our data reveals a crucial difference in colour preference between Australian (*A. mjoebergi*) and American (*L. leptodactyla*) fiddler crabs. In the absence of ultraviolet information, while female Australian fiddler crabs are attracted by enlarged yellow (Detto et al. 2006; Dyson et al. 2020) and orange claws (Dyson et al. 2020), female American fiddler crabs present no sensory bias to any hypertrophied claw colouration whatsoever, as shown by our second experiment.

### The Role of Claw Colouration in Fiddler Crabs The private communication channel hypothesis

On one hand, our visual modelling results do not support the existence of a private communication channel (Cummings et al. 2003), through which fiddler crabs could communicate without being seen by avian predators. In fact, our data show the opposite of what we predicted in our first hypothesis. Birds, which are responsible for most predation pressure suffered by fiddler crabs (Ribeiro et al. 2019), should outperform female fiddler crabs in using colour to identify males’ hypertrophied claws on the sandy substrate. These results are in line with what has been reported by previous visual modelling studies conducted in other species of fiddler crabs (Hemmi et al. 2006; Cummings et al. 2008).

### The handicap principle hypothesis

On the other hand, when considering our behavioural and visual modelling data, both seem to corroborate our second and third hypotheses. Previous allegations that male hypertrophied chelae would assist fiddler crab detection by humans (Jordão and Oliveira 2001), in addition to our modelling results predicting that birds should pose a threat to male fiddler crabs, demonstrate that males’ hypertrophied claws might be regarded as handicaps (Zahavi 1975). According to Zahavi’s point of view (Zahavi and Zahavi 1997), the simple act of bearing a handicap (i.e., flamboyant claw) and not having been captured by predators would prove an individual’s quality to their potential mates, as they propose it would happen with peacocks and other species displaying bright colourations. So, we could speculate that just a few high-quality males should be able to break claw crypticity and pay the costs for socio-sexual advertisement (i.e., honest signal), coping with the resulting enhancement in bird predation pressure.

Another, not mutually exclusive, possibility would be that colourful chelipeds could serve as anti-predatory honest signalling. In an exquisite behavioural study, Bildstein et al (1989) showed that the enlarged claw of male fiddler crabs reduces the likelihood of their capture by relatively large avian predators (i.e., white ibises), enhancing the chances that declawed males, or females, have of being chosen instead. These results are also in line with Zahavi’s handicap principle, since white ibises would be choosing to attack prey that could not convey reliable proofs of their quality (i.e., fiddle crabs with small, or absent, chelae), in the same way wolves prefer to attack gazelles that run instead of jumping (Zahavi and Zahavi 1997).

### Fisherian hypotheses

Female choice in fiddler crabs might be linked to direct (i.e., resources, protection) and indirect (i.e., better sperm, good genes) benefits that are supplied by the males. A good example of direct benefits, that might be explored by females, is the existing relation between size of hypertrophied claws and quality of male burrows, with larger males occupying larger, safer, and more thermally stable burrows (Christy 1987). Concerning the potential indirect benefits of female choice, two dominant hypotheses have been recognized, Fisher’s runaway selection hypothesis and the good-genes hypothesis (Anderson 1994).

Although our study has not evaluated if there is a genetic correlation between UV reflection by male claws and females’ preference for UV cues, another way of interpreting our data would be that a Fisherian runaway selection is under action in fiddler crabs, selecting a strong female preference for UV colouration in correlation with a strong UV reflection by males’ hypertrophied claws. According to the runaway hypotheses, in which a self-reinforcing process of ever-elaborating traits and preferences would take place, the mean values of traits and preferences would increase, while their variances and correlation should approach a stable equilibrium (Henshaw and Jones 2020). Our data, however, does not seem to support Detto and Backwell (2009) Fisherian hypothesis, in which selection for UV reflection could be explained by the enhancement of male conspicuity with female concomitant attraction. Even though we have shown there to be a clear preference for UV signals among half of female fiddler crab population, our visual models predict that only the claws of a very few males (i.e., 17.5% of the population) would be conspicuous to their mates, whilst the remaining of them would be regarded as cryptic, something that goes against the logic of Fisher’s classical runaway selection hypothesis.

Nevertheless, a recent reinterpretation of Fisher’s original mathematical model predicts the occurrence of two qualitatively different outcomes besides the classical runaway: explosive runaway and fizzle away (Henshaw and Jones 2020).

According to the explosive runaway possibility, in case a large variance in preference overcame fiddler crab’s female population, with some females showing a huge preference for male UV reflection while others could not care less about the same male trait, selection could be so strong that extreme outliers (i.e., those few males with conspicuous claws) in the original distributions could be strongly favoured, leading to a super exponentially increase in both the means and variances of traits and preferences, that could reach absurd values in very few generations. When we take the behavioural data from our first experiment and split every female into two categories, we can see that half of them choose an UV-reflecting male (n = 27), while the other half does not (n = 33). If Henshaw and Jones (2020) correction of Fisher’s mathematical model is accurate, this observed variance in female’s preference for UV signals should start an ultra-rapid selection process with explosive increase in enlarged claws UV conspicuity, something that we also have not encountered in our sample.

At last, we conclude that the third Fisherian possible outcome, the fizzle away selection, in which variation in both traits and preferences would converge to zero, while the means of both traits and preferences would plateau after an initial period of increase (Henshaw and Jones 2020), also does not fit our results, inasmuch as we have shown variation to take place both on female preference for UV signals and UV reflectance by male enlarged claws.

Only further experimentation testing, for example, if there is a hereditarian cooccurrence between a higher UV reflection in males and a higher UV preference in related females, or how physiological attributes of male fiddler crabs correlate with their colour signals, especially if female choice leads to offspring of superior viability, can give us more conclusive answers about the evolution of UV reflection on fiddler crabs in light of Fisher’s and the good-genes hypotheses.

## Conclusions

We failed to prove the existence of a UV private communication channel related to fiddler crabs’ enlarged claw colouration. Our data endorse the view that male enlarged UV-reflecting chelae function as handicaps and might honestly signal quality to potential mates and predators. We also have shown that, similarly to Australian fiddler crabs, American fiddler crabs produce, discriminate, and prefer UV light signals. Would female preference for UV signals and UV reflection by male enlarged claws be widespread traits within fiddler crabs? Have the trait and the preference evolved only once, a long time ago, before the split of Gelasiminae into the Indo-West Pacific and American groups (Shih et al. 2016)? Or, instead, are they more recent acquisitions that have been independently selected in the American and Indo-West Pacific branches through convergent evolution? Future studies investigating traits and preferences in additional species of fiddler crabs can shed more light on how the evolution of UV signalling happened in this group.

## ACKNOWLEDGMENT

We are deeply indebted to J. Medeiros and L. V. Silva, for helping with the logistics of the project. We would also like to thank H. R. Silva, M. S. Capitulo, J. B. C. Neto and B. A. Souza for assistance in data collection, Prof. Glenn W. Erickson (English native speaker) for reviewing our manuscript, and Prof. Natan Hart for sharing *Pavo cristatus* ocular transmission spectrum. This study was financed in part by the Coordenacao de Aperfeicoamento de Pessoal de Nivel Superior – Brasil CAPES (Finance Codes 001 and 043/2012), by Conselho Nacional de Desenvolvimento Cientifico e Tecnologico – Brasil (CNPq) (Finance Codes 478222/2006-8 and 474392/2013-9) and by Programa de Apoio aos Nucleos de Excelencia – FAPERN/CNPq (Finance Code 25674/2009). An Undergraduate Scientific Initiation Scholarship was granted to R.S.G., M.Sc. Scholarships were granted to D.J.A.S. and M.F.E., and a Researcher Scholarship was granted to D.M.A.P., all by Conselho Nacional de Desenvolvimento Cientifico e Tecnologico – Brasil CNPq.

## APPENDIX

**Appendix 1.**
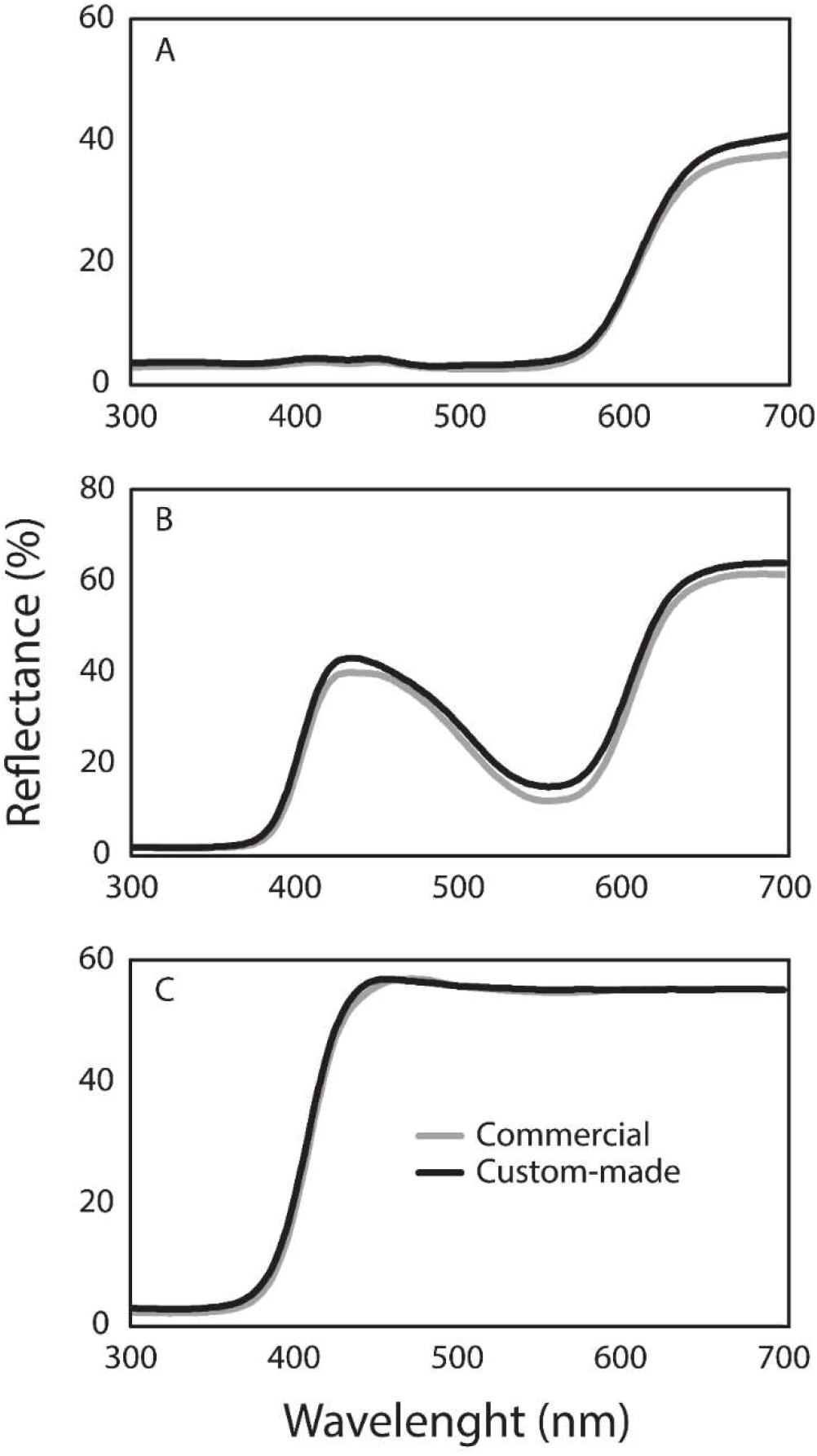
Reflectance spectra, of three different natural surfaces, measured with aid of either a commercial reflectance probe holder (RPH-1, Ocean Optics Inc., Dunedin, Florida) or a custom-made, 3D printed, reflectance probe holder. A) Petal of *Delonix regia* (Bojer ex Hook.) Raf.; B) Petal of *Catharanthus roseus* (L.) G.Don; C) Petal of *Plumeria pudica* Jacq.

